# Modeling features of addiction with an oral oxycodone self-administration paradigm

**DOI:** 10.1101/2021.02.08.430180

**Authors:** Caitlin A. Murphy, Yu-Hsuan Chang, Rajesh Pareta, Jun-Nan Li, Richard A. Slivicki, Tom Earnest, Jessica Tooley, Kavitha Abiraman, Yvan M. Vachez, Robert W. Gereau, Bryan A. Copits, Alexxai V. Kravitz, Meaghan C. Creed

## Abstract

Prescription opioid use is an initiating factor driving the current opioid epidemic. There are several challenges with modeling prescription opioid addiction. First, prescription opioids such as oxycodone are orally self-administered and have different pharmacokinetics and dynamics than morphine or fentanyl. This oral route of administration determines the pharmacokinetic profile, which is critical for establishing reliable drug-reinforcement associations in animals. Moreover, the pattern of intake and environment in which addictive drugs are self-administered intake are critical determinants of the levels of drug intake, sensitization and relapse behavior. This is an important consideration with prescription opioid use, which is characterized by continuous drug access in familiar environments. Thus, to model features of prescription opioid use and the transition to abuse, we present an oral oxycodone self-administration paradigm that is administered in the home cage. Mice voluntarily self-administer oxycodone in this paradigm without any taste modification such as sweeteners, and exhibit preference for oxycodone, escalation of intake, physical signs of dependence, reinstatement of seeking after withdrawal, and a subset of animals demonstrate drug taking that is resistant to aversive consequences. This model could be useful for studying the neurobiological substrates specifically relevant to prescription opioid abuse.

## Introduction

Misuse of prescription opioids is a significant public health and economic burden worldwide. In 2017 in the United States, more than 10 million people reported prescription opioid misuse [1]. Despite its importance, modeling prescription opioid abuse in rodents has been challenging [2–5]. Certain aspects unique to prescription opioid use, such as route of administration and pattern of intake, may contribute to abuse liability and propensity toward relapse. As such, a representative preclinical model of prescription opioid addiction is critical for understanding the neurobiology underlying this disorder.

Pre-clinical drug self-administration studies have typically employed paradigms where drug administration is limited to discrete time periods in operant chambers, before returning animals to their home cages. In these studies, drugs are often self-administered intravenously under various reinforcement schedules, which allows experimenters to assay distinct features of addiction behavior, such as motivation for the drug [6,7], reinstatement of drug seeking [8,9] or compulsive drug taking [10–12]. However, several characteristics of prescription opioid use introduce unique challenges that may not be optimally modelled by classical self-administration paradigms.

First, in contrast to many addictive drugs (e.g., heroin, cocaine, methamphetamine, or morphine), prescription opioids are most often orally self-administered: among chronic and recreational opioid users, oral self-administration is decidedly preferred over non-oral routes (e.g., insufflation, injection) [13,14]. The route of administration of drugs of abuse is highly relevant for establishing learned drug associations [15], which contribute to the subjective reinforcing properties of the drug, escalation of intake, and propensity toward reinstatement [10,11,16–20]. Indeed, the pharmacodynamic and pharmacokinetic profiles of opioids – and therefore their physiology and time-course of action – vary widely across routes of administration [21]. These differences underscore the need to model prescription opioid abuse through the oral route of administration.

Second, prescription opioids are most commonly self-administered in a familiar setting such as the home (see Caprioli et al. (2007) for a review [22]). Yet, where oral self-administration has been reported [2–4], experimental paradigms have required self-administration to take place in a novel context (e.g., operant chamber) and under food restriction. Environmental context (i.e., setting of drug taking) plays an important role in both drug taking and reinstatement, a finding that has been well-documented in humans and recapitulated by animal models [22–27]. Moreover, the nature of operant self-administration paradigms requires intermittent access to the drug during daily sessions separated by acute periods of forced abstinence. However, this intermittent pattern of intake stands in stark contrast to continuous access to prescription opiates, which is even more pronounced in the case of extended-release formulations [28]. The pattern of drug intake is critical, since intermittent access patterns exacerbates mesolimbic adaptations induced by opioid self-administration [29,30].

Previous investigations have applied the oral route of self-administration by implementing two-bottle choice procedures, where animals are free to choose between bottles containing drug- and drug-free solutions. Because the opioid alkaloid structure confers a bitter taste to opium derivatives, and typically sweeteners have been added to encourage animals to self-administer opioid via the oral route, or adulterants to the drug-free bottle to control for aversive taste. However, the addition of these extraneous reinforcers or adulterants introduce experimental confounds which limit the interpretability of two-bottle choice procedures [31]. For example, sucrose itself can support seeking after abstinence [32], disrupt overall patterns of fluid intake [33], and stimulate dopamine and opioid receptors in the mesolimbic pathway [34–37]. Therefore, recapitulating a continuous access pattern and oral route of administration without adulterants is important to accurately model prescription opioid use disorder (OUD).

Here, we characterize a home cage-based, oral oxycodone (OxyContin, Tylox^TM^ or Percodan^TM^) self-administration paradigm that models several features of prescription opioid abuse. We demonstrate that mice exhibit a preference for high concentrations of unadulterated oxycodone (1.0 mg/mL) over regular drinking water and voluntarily escalate drug intake over the course of a two-bottle choice paradigm. This model also recapitulates multiple behavioral features that have been used to model criteria for addiction as defined by the fifth version of the diagnostic and statistics manual [40] (DSM), specifically escalation of drug intake, physical signs of dependence, drug craving after withdrawal and drug use despite negative consequences. Critically, these behaviors were also accompanied by potentiation of excitatory synapses in the nucleus accumbens (NAc), which is a conserved neurobiological substrate of drug reinstatement [41–43]. By understanding the neurobiological substrates specific to prescription opioid abuse, it may be possible to design effective therapies for prescription opioid misuse, and to develop novel medications to treat pain with lower abuse liability.

## Materials and Methods

### Subjects

A total of 112 adult mice (C57BL/6J background; age, 10–16 weeks). Both males (n=57) and females (n=55) were included in the study and balanced across experiments and treatment groups. Mice were housed in standard mouse vivarium caging and kept on a 12 h light/dark cycle (light onset, 7:00 A.M.; light offset, 7:00 P.M., temperature (22-26°C), in-cage humidity (22-50%). Mice had ad libitum access to food and were single housed during self-administration experiments. Experiments were run in successive waves of treatment- and sex-matched cohorts with up to 20 mice being run simultaneously. All studies were approved by the internal animal care and use committee at Washington University in St. Louis.

### Home cage lickometer devices

All devices used in these experiments are low-cost and open-source, with detailed parts lists, code and fabrication instructions published and available on-line [44,45]. Drinking water and oxycodone solutions were provided using an in-cage two-bottle lickometer apparatus which detects interactions with two separate drinking spouts via a pair of photobeams [45]. Cages were also equipped with a wireless passive infrared (PIR) activity monitor [44] to measure mouse activity levels (Models 4430 and 4610 from MCCI Corporation, Ithaca NY). Lickometer counts on each sipper, PIR activity data, and environmental measurements (temp, humidity, light levels) were transmitted via Low frequency Radio Wide Area Network (LoRaWAN) using Internet of Things (IoT) infrastructure (The Things Network, Netherlands), saved in a cloud-database (InfluxDB), and visualized with an online dashboard (Grafana). The wireless lickometer and PIR sensors were also equipped with back-up microSD cards and liquid volumes were measured manually each day.

### Oxycodone self-administration

<u>Habituation:</u> Mice were transferred to single housing with an in-cage two-bottle choice apparatus at least three days prior to the start of the escalation protocol, during which the sipper device was the only source of drinking water (supplied in both tubes). <u>Phase I:</u> Mice in the experimental group (“OXY mice”) were supplied with a single bottle of oxycodone hydrochloride dissolved in drinking water as their only source of liquid; 0.1 mg/mL was available for 24 hours, 0.3 mg/mL was available for 48 hours, and 0.5 mg/mL was available for 48 hours (Fig 1A), based on an operant protocol developed by Phillips et al. [3]. Mice in the control group (“CTRL mice”) had access to drinking water only throughout the protocol. The position of the bottle within the two-bottle apparatus was switched daily. <u>Phase II:</u> Oxycodone (1.0 mg/mL in drinking water) and drinking water were supplied to OXY mice ad libitum in the two-bottle choice apparatus. Both solutions were available 24 hours a day for 7 days. Drinking water was supplied in both bottles for CTRL mice. <u>Withdrawal:</u> The two-bottle lickometer was removed from the home cage after the 7-day two-bottle choice period, and standard cage-top water bottles were replaced as the source of drinking water. Separate groups of mice were used for the oxycodone seeking, quinine adulteration and electrophysiology studies outlined below.

**Figure 1.**
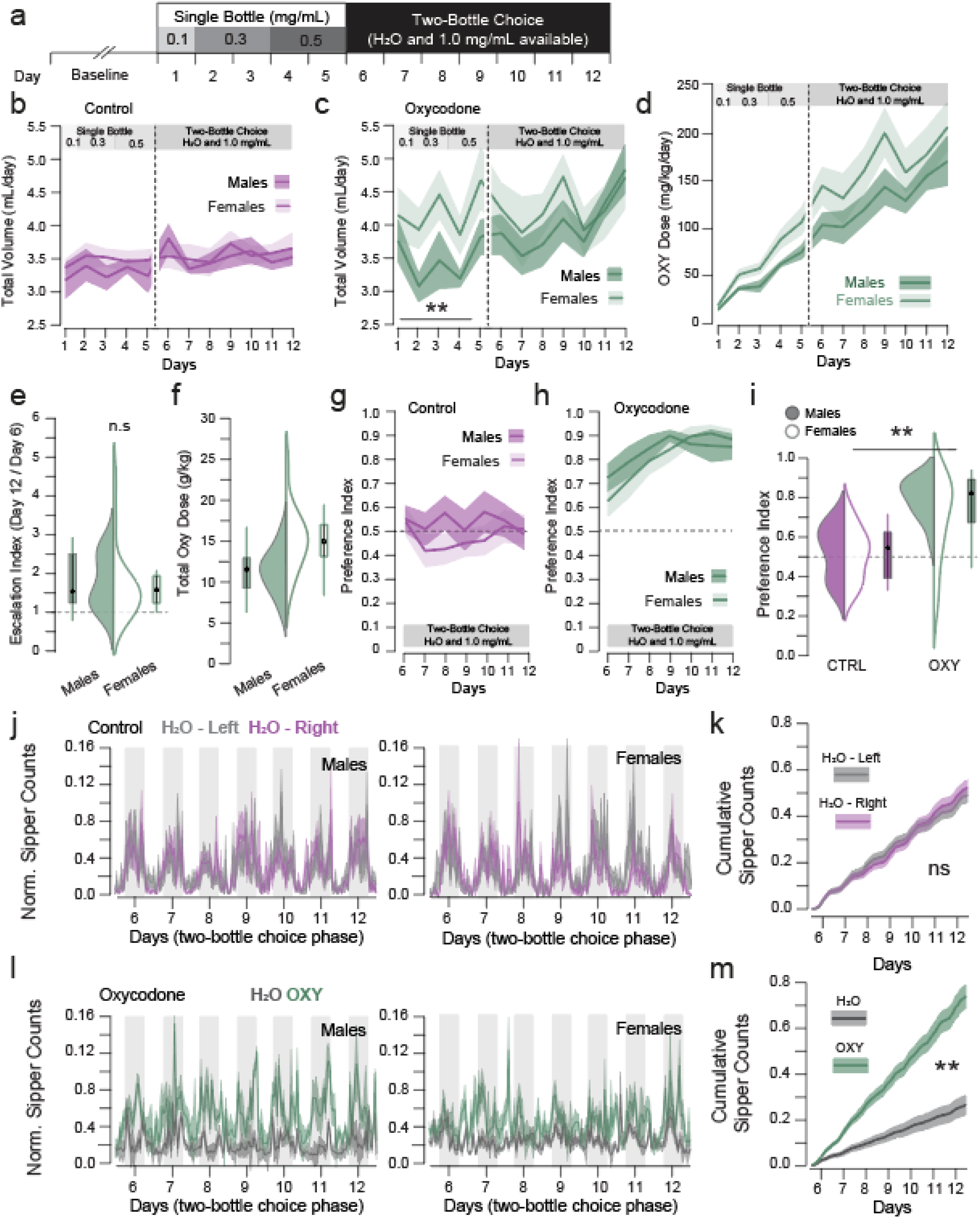
Mice voluntarily consume oxycodone in a two-bottle choice paradigm. **(a)** schematic of experimental paradigm (n = CTRL: 20M/20F, OXY: 26M/26F). **(b-c)** Volumes of total liquid consumed in control or oxycodone-administering mice (CTRL female 3.527 ± 0.044, male = 3.422 ± 0.049, OXY female = 4.233 ± 0.103, male = 3.862 ± 0.083). Mice with OXY available drank significantly higher volumes over the two-bottle choice phase (F_drug_ = 55.679, p < 0.001), and female mice consumed higher volumes of OXY during the self-administration protocol (F_sex_ = 9.588, p = 0.002, F_sex●drug_ = 1.833, p = 0.045). **(d)** Oxycodone dosage increased throughout the single bottle phase and mice continued to escalate their consumed oxycodone levels throughout the two-bottle choice phase (F_Time_ = 2.743, p = 0.002). **(e)** There was no sex difference in escalation index, calculated as the dose on the final day of two-bottle choice (day 12) divided by dose on the first day of the two-bottle choice phase (day 6; Females = 1.610 ± 0.193, Males = 1.748 ± 0.153, F = 0.313, p = 0.579). **(f)** Female mice self-administered higher total doses of oxycodone over the course of the protocol (Females = 1499.135 ± 78.745, Males = 1147.788 ± 59.855, F = 12.619, p = 0.001). **(g-h)** OXY-self-administering mice, but not control mice developed a significant preference for the oxycodone-containing bottle over the 7 days of two-bottle choice, which did not differ between males and females (F_drug_ = 227.488 p < 0.001, F_drug●day_ = 9.233, p = 0.003, F_sex_ = 3.308, p = 0.070). **(i)** There was no systematic preference for drinking water from the left or right bottle in CTRL mice, while mice consumed significantly more OXY solution relative to drinking water when available (CTRL female = 44.8 ± 4.9%, male = 53.9 ± 6.0% preference for right drinking bottle; OXY female = 88.2 ± 3.9%, male = 89.6 ± 3.2% preference for OXY-containing bottle, F_drug_ = 76.45 p < 0.001). **(j-k)** Sipper counts (photobeam breaks on each lickometer) were normalized for total Sipper Counts made over the 7 days of the two-bottle choice task and plotted in 1 hour bins in CTRL mice. There was no difference in sipper counts on the left vs. right sipper in control mice (CTRL female Left/Right = 0.531 ± 0.039/0.466 ± 0.039, male Left/Right = 0.465 ± 0.061/0.471 ± 0.048). **(l-m)** Counts were significantly higher on the oxycodone-containing bottle relative to the water containing bottle in oxycodone-administering mice (OXY female H_2_O/OXY = 0.287 ± 0.075/0.641 ± 0.087, male H_2_O/OXY = 0.218 ± 0.036/0.782 ± 0.036, F_bottle_ = 29.485, p < 0.001, F_drug●bottle_ = 38.154, p < 0.001).

### Oxycodone Seeking Test

In a subset of mice (n_CTRL_ =10M/10F, n_OXY_ = 10M/10F), an oxycodone seeking test was performed. After 24 hours (early) or 22-24 days (late) withdrawal, the lickometer device containing 2 empty bottles was re-introduced to the cage for a 60-minute probe trial, and interactions with the sipper tubes as detected by the photobeam were recorded. Mice were tested at both early and late withdrawal points.

### Quinine Adulteration

In a subset of mice (n_CTRL_ = 8M/7F, n_OXY_ = 8M/7F) we tested resistance of oxycodone drinking to quinine adulteration. Following the conclusion of the 7 days of two-bottle choice protocol, increasing concentrations of quinine were added to the oxycodone-containing bottle (OXY mice), such that OXY mice had the choice between standard drinking water (one bottle) and 1.0 mg/mL oxycodone + quinine. CTRL mice had the choice between standard drinking water (one bottle) and water + quinine. Mice were given 48 hours at each quinine concentration in increasing order: 125 µM, 250 µM, 325 µM, volumes consumed from each tube were recorded daily.

### Baseline Oxycodone Preference Test

In a separate cohort of naïve mice (n = 19M/22F), we determine baseline preference for oxycodone by giving the mice the choice between 1 bottle of drinking water and 1 bottle of 1.0 mg/mL oxycodone hydrochloride solution. Preference was calculated as the proportion of oxycodone as a fraction of total liquid volume consumed over the overnight session.

### Physical Withdrawal Signs

<u>Acquisition:</u> After either 24 hours or 22-24 days of withdrawal, mice were habituated to the testing room for at least 1 hour. Three consecutive, 5 minute videos were acquired of individual mice at 100 fps in a double-tall Von Frey chamber with an unobstructed background and overhead lighting (LUX). Videos were scored for signs of withdrawal (jumps, tremors, movement speed) and anxiety related behaviors (grooming, rearing, climbing) using markerless pose estimation and subsequent supervised machine learning predictive classifiers of behavior. Markerless pose estimation was done using DeepLabCut^TM^ (DLC, version 2.2b6 [46,47]): 12 frames of 64 videos representing all treatment conditions were extracted using k-means clustering and subsequently manually annotated for the following 9 body parts: left ear, right ear, left forepaw, right forepaw, left hind paw, right hind paw, snout, tail base, back. The training fraction was set to 0.95, and the resnet_50 network was trained for 1,030,000 iterations. A train error of 1.82 and test error of 11.47 were achieved with a cutoff value of p=0.6. DLC tracking data and generated videos were then imported to the Simple Behavioral Analysis (SimBA) project workflow (version 1.2 [48]). Within SimBA, single animal, 9-body part supplied configuration was used to extract behavioral features from the pose estimation data after outlier and movement correction (both parameters set to 7x outside the interaural distance. Extracted frames from four independent videos were annotated to build classifiers for the following behaviors: “climbing”, “jumping”, “tremor”, “rearing, and “grooming”. Individual models were trained using a random forest machine model with 2000 estimators, and a training fraction of 0.2 (default hyperparameters). Following validation, videos were analyzed using the random forest model, with the following p-cutoffs and minimum behavioral bout lengths for each of the following behaviors: “climbing (p = 0.26, 35 ms)”, “jumping (p = 0.4, 35 ms)”, “tremor (p=0.0495, 50 ms”, “rearing (p = 0.45, 70 ms)”, and “grooming (p = 0.38, 70 ms)”. The SimBA pipeline is built primarily on scikit-learn [49], OpenCV [50], FFmpeg45 [51], and imblearn [52]. Total time spent in each behavior was exported and the three videos were averaged to get one value for each subject to be included in the analysis. Euclidean distance of displacement of all body parts was also extracted from the pose-estimation data and averaged to achieve one ‘movement’ score for each subject.

### Patch Clamp Electrophysiology

Coronal mouse brain slices, 220 μm in thickness were prepared in cooled artificial cerebrospinal fluid containing (in mM): 119 NaCl, 2.5 KCl, 1.3 MgCl, 2.5 CaCl_2_, 1.0 Na_2_HPO_4_, NaHCO_3_ 26.2 and glucose 11, bubbled with 95% O_2_ and 5% CO_2_. Slices were kept at 30-32°C in a recording chamber perfused with 2.5 mL/min artificial cerebrospinal fluid. Visualized whole-cell voltage-clamp recording techniques were used to measure spontaneous and synaptic responses of NAc shell MSNs. Holding potential was maintained at −70 mV, and access resistance was monitored by a depolarizing step of −10 mV each trial, every 10 s. Currents were amplified, filtered at 2 kHz and digitized at 10 kHz. All experiments were performed in the presence of picrotoxin (100 μM) to isolate excitatory transmission, and TTX (0.5 µM) was included for recordings of miniature excitatory post-synaptic currents (mEPSCs). Internal solution contained (in mM): 130 CsCl, 4 NaCl, 5 creatine phosphate, 2 MgCl_2_, 2 Na_2_ATP, 0.6 Na_3_ GTP, 1.1 EGTA, 5 HEPES and 0.1 mm spermine. Synaptic currents were electrically evoked by delivering stimuli (50 – 100 μs) every 10 seconds through bipolar stainless-steel electrodes. The AMPAR component was calculated as the peak amplitude at −70 mV, The NMDAR component was estimated as the amplitude of the outward current at +40 mV after decay of the AMPA current, measured 50 msec after the electrical stimuli (Fig 5A-C). Paired-pulse ratio PPR was calculated as the ratio of the second to first baseline-subtracted peak elicited with an ISI of 50 msec.

### Statistical Analyses

Photobeam break data on the lickometer devices was extracted and analyzed using custom python code (SipperViz graphical user interface is available for download at https://github.com/earnestt1234/SipperViz). Statistics were performed in python (3.7 using Spyder 4.1.5), using the pinguoin (0.3.10) and statsmodels (0.10.0) packages. Data was analyzed with repeated measures ANOVA or two-factor ANOVA where appropriate, followed by post-hoc t-tests. Sex and Treatment group were included as between-subject factors for all analyses.

## Results

### Mice exhibit preference, escalation, and altered pattern of intake of oxycodone

We mimicked a prescribed course of opioids by including escalating concentrations of oxycodone in the drinking water in a single-bottle phase, followed by a two-bottle choice phase where mice could choose between a concentrated oxycodone solution (1.0 mg/mL) and unadulterated drinking water (Fig 1A-C). During the single bottle phase, mice increased their daily oxycodone consumption (Fig 1D), which is predicted, based on the increasing concentrations of oxycodone in their drinking water (0.1, 0.3, 0.5 mg/mL). However, mice continued to systematically escalate their drug intake during the two-bottle choice phase, where the drug concentration remained constant (1.0 mg/mL) and drinking water was also available ad libitum (Fig 1D-F), with OXY mice consuming higher total volumes of liquid than CTRL mice that only had access to drinking water (Fig 1B,C). Moreover, during this two-bottle choice phase, mice exhibited a significant *preference* for 1.0 mg/mL oxycodone over drinking water (Fig 1G-I), which is noteworthy, since 1.0 mg/mL oxycodone solution is bitter, and both male and female naïve mice preferred drinking water when given the choice between drinking water and 1.0 mg/mL oxycodone (Fig S1A).

In addition to the total volume consumed, the home cage sipper device allows for 10 second resolution of licking behavior on each bottle. Examination of the time course of the drinking behavior confirmed that there was no systematic preference for the left or right bottle in CTRL mice (Fig 1J-K), while OXY mice exhibited higher levels of responding on the oxycodone-containing bottle (Fig 1L-M). Moreover, the circadian index of sipping behavior (defined as sipper counts registered during the dark cycle / sipper counts registered during the light cycle) was significantly reduced in OXY mice relative to controls (Fig S3). The home cage nature of this paradigm allows for large sample sizes with statistical power to detect sex differences (n_CTRL_=40 and n_OXY_=52). While both male (n=26) and female (n=26) mice exhibited preference for oxycodone (Fig 1H-M, S1C, D), during the single bottle phase, female mice consumed more oxycodone than males (Fig 1C). Female mice continued to self-administer higher doses of oxycodone over the two-bottle phase of the protocol (Fig 1D, S1E-F). No sex differences were observed for escalation index or oxycodone preference (Fig 1G-M, S1C,D, S2).

### Oxycodone self-administration induces physical signs of intoxication and withdrawal

DSM criteria for substance use disorder include tolerance to and withdrawal from an addictive substance. Although we did not measure drug tolerance, the oral self-administration protocol did induce physical signs of intoxication (hyperactivity) as well as acute and protracted physical withdrawal symptoms (Fig 2). Acute injections of oxycodone [53–55], or vapor exposure [56] to opioids induce hyperactivity, and these locomotion-inducing properties of addictive drugs are thought to reflect increased dopamine release in the NAc [57–59]. We equipped our home cages with PIR sensors to detect locomotor activity during the self-administration protocol (Fig 2A). OXY mice exhibited higher levels of hyperactivity than CTRL mice during both the single- and two-bottle choice phases of self-administration (Fig 2B,C), suggesting that levels of oxycodone voluntarily self-administered via the oral route produced sufficient brain levels of oxycodone to induce hyperactivity [54,60,61]. We found no sex differences in oxycodone-induced locomotor activity (Fig S4).

**Figure 2.**
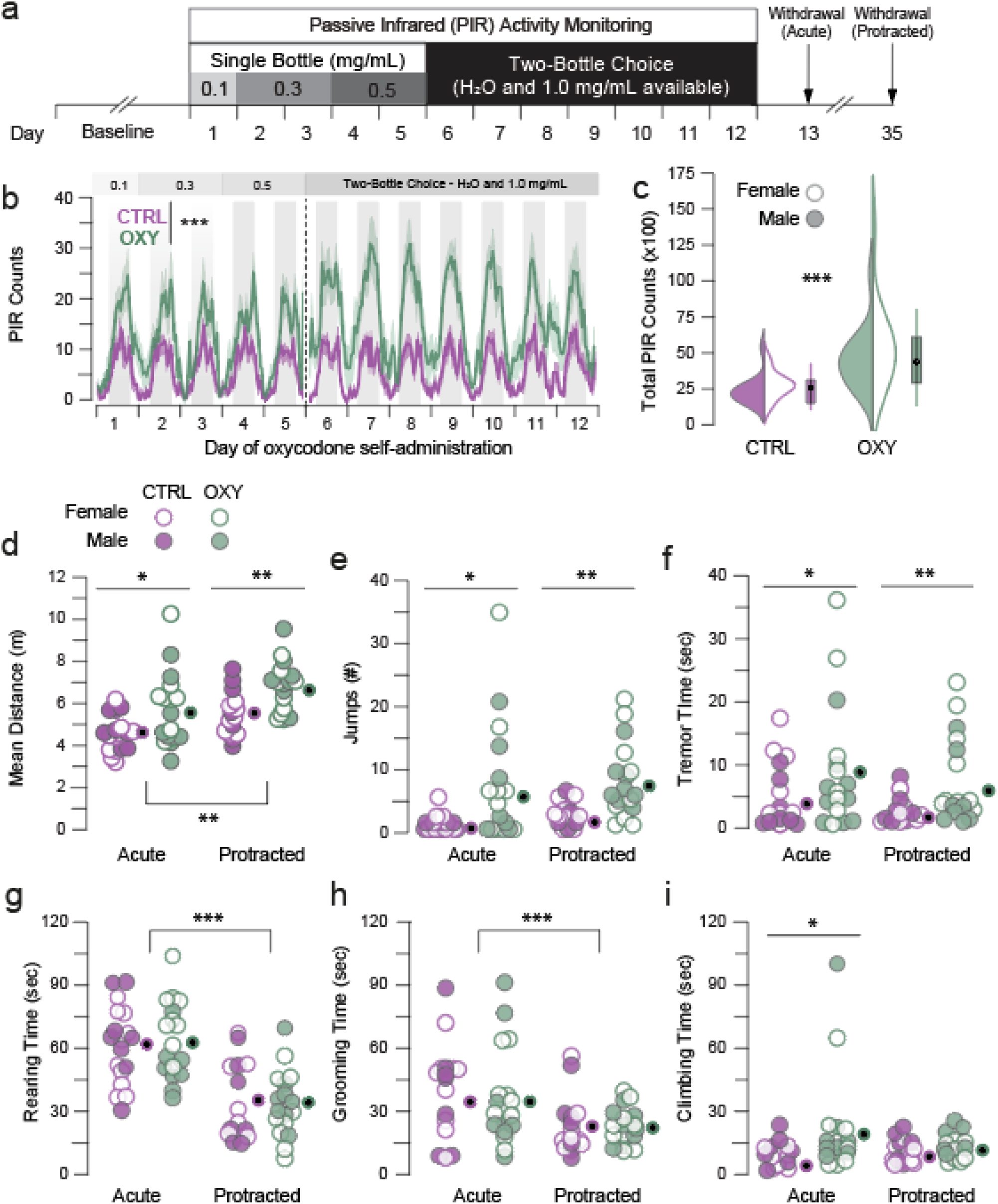
Oral oxycodone self-administration induces hyperactivity and persistent physical withdrawal signs. **(a)** Experimental schematic. Home cages were equipped with PIR sensors to monitor activity during the single- and two-bottle choice phases of OXY-self administration. Withdrawal symptoms were assayed after 24h and after 3 weeks of withdrawal. **(b-c)** Both male and female mice exhibited significantly higher homecage activity during OXY self-administration than control mice (CTRL: 2072.189 ± 196.455, OXY = 4530.370, F_sex_ = 0.956, p = 0.333, F_drug_=19.549, p<0.001). **(d)** There was a significant effect of drug and time point, with OXY mice exhibiting higher levels of movement relative to control mice at acute (CTRL: 473.18 ± 20.18, OXY: 570.65 ± 43.45 cm) and protracted (CTRL: 555.56 ± 27.62, OXY: 689.67 ± 29.00 cm) withdrawal time points (F_TimePoint_=10.938, p = 0.002, F_drug_ = 13.343, p=0.001). **(e)** OXY mice exhibited greater numbers of jumps at both early (CTRL: 0.778 ± 0.298, OXY: 6.824 ± 2.254) and late withdrawal points (CTRL: 2.400 ± 0.498, OXY: 7.735 ± 1.523, F_drug_ = 15.284, p < 0.001). **(f)** OXY mice exhibited increased paw tremors at both early (CTRL: 4.053 ± 1.217, OXY: 9.132 ± 2.423) and late time points (CTRL: 2.420 ± 0.590, OXY: 6.025 ± 1.763, F_drug_ = 7.155, p = 0.010). **(g-h)** While there was a significant effect of time point on both rearing and grooming behavior (Rearing F_TimePoint_ = 46.380, p < 0.001, Grooming F_TimePoint_ = 7.757, p = 0.007), these behaviors were not different between control and OXY-self-administering mice. **(i)** There was a significant effect of drug on climbing time at the acute time point only (CTRL: 7.862 ± 1.269, OXY: 21.092 ± 6.080 sec, F_drug_ = 5.065, p =0.028). (h-i) *p<0.05, **p<0.01, ***p<0.001.

We also determined whether withdrawal from oral oxycodone self-administration induced physical withdrawal signs. During withdrawal from opioids such as morphine or heroin, mice exhibit characteristic withdrawal signs including tremors, jumping, teeth chattering and diarrhea. We quantified jumping and tremor, along with the total distance moved by the animal. Relative to CTRL mice, OXY mice exhibited significantly more movement (Fig 2D), as well as increased jumping (Fig 2E) and paw tremors (Fig 2F) at both the acute (1 day) and protracted (3 weeks) withdrawal periods.

Finally, we quantified climbing (an index of escape behavior), grooming, and rearing, which have been used as a proxy for anxiety-like behavior, and have also been reported to be increased following withdrawal from opioids. OXY mice did not exhibit significantly more rearing or grooming relative to CTRL mice (Fig 2G, H). While there was a significant effect of drug on escape behaviors in acute oxycodone withdrawal, this effect was driven by two subjects and resulting group differences were small (Fig 2I). There were no significant differences between male and female mice for any physical symptom measured. Together, these results demonstrate that oral oxycodone self-administration induces physical signs of dependence that are consistent with opioid withdrawal syndrome.

### The two-bottle choice paradigm induces aversion-resistant drug consumption and persistent drug seeking into withdrawal

Drug seeking behavior is operationally defined as performance of an action that previously led to consumption of the drug, in the absence of the drug itself [62,63]. In the case of i.v. drug self-administration, this response may be operationalized as lever-pressing or nose-poking to receive a drug infusion. In our model, oxycodone is delivered through a unique sipper device, which is distinct from the standard cage-top water bottle. Therefore, we assayed drug seeking behavior by measuring photobeam breaks on the lickometer devices under extinction conditions, with no oxycodone or drinking water in the tubes after 24 hours and 3 weeks of forced abstinence from oxycodone (Fig 3A). OXY mice exhibited significantly higher lickometer counts than CTRL mice in both the acute (1 day) and protracted (3 weeks) withdrawal periods (Fig 3B). This drug seeking behavior was not significantly different between the early and the late probe trials, nor between males and females (Fig 3C-H, Fig S5A-B). This establishes that the two-bottle choice paradigm supports drug seeking behavior that persists after abstinence.

**Figure 3.**
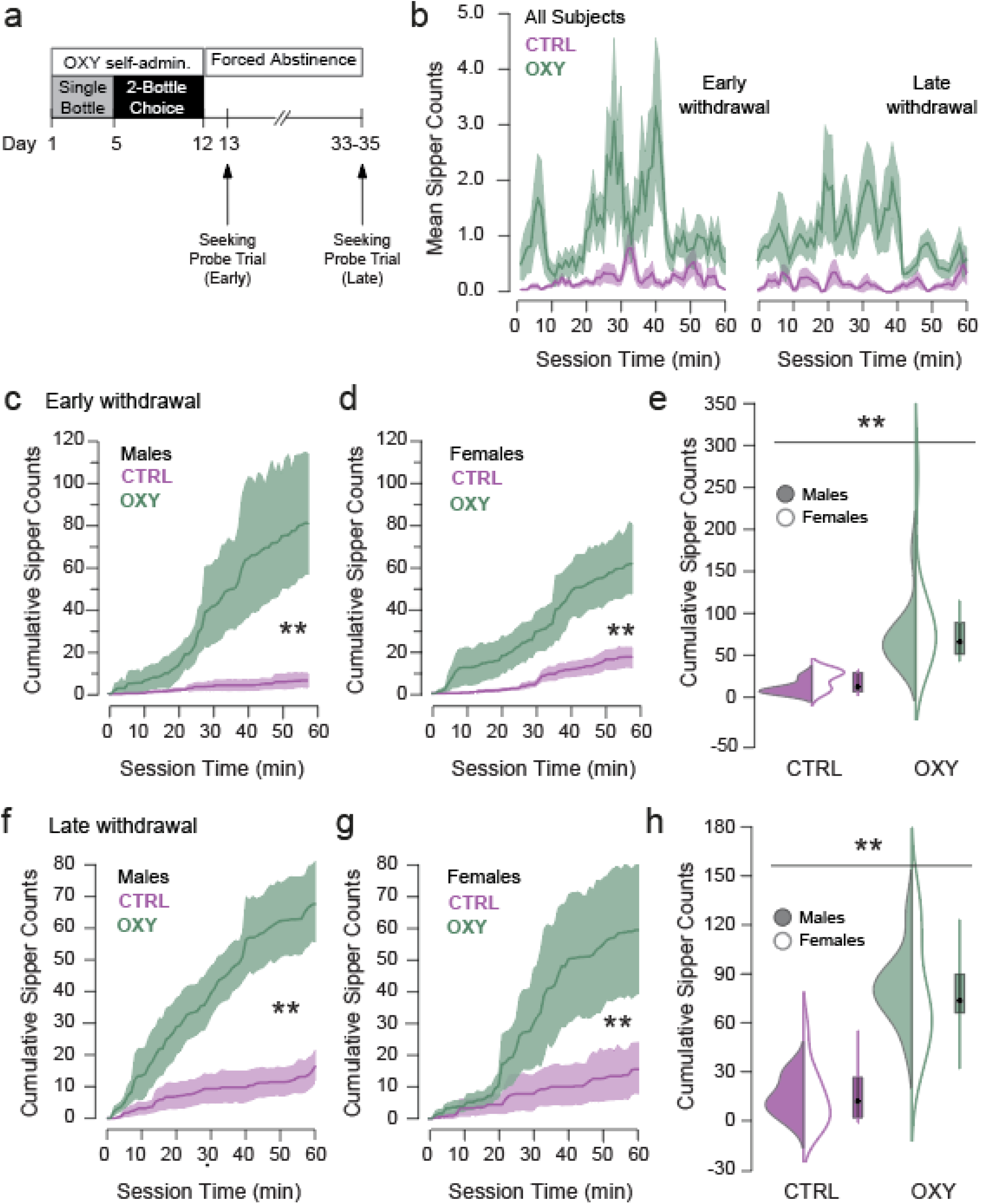
Drug seeking behavior persists into protracted withdrawal. **(a)** Experimental timeline (CTRL n = 10M/10F, OXY n = 10M/10F). 24h (early seeking) and 24 days (late seeking) after the two-bottle choice protocol, drinking counts were measured under extinction conditions. **(b)** Time course of photobeam breaks (sipper counts) made on empty bottles over the 60 minute probe trial; mice that had self-administered OXY showed more seeking interactions relative to controls (F_drug_ = 51.062 p < 0.0001), with no difference between early and late seeking (F_TimePoint_ = 0.227, p = 0.653, F_drug●TimePoint_ = 0.927, p=0.339). **(c-e)** Both male and female mice that had self-administered OXY exhibited higher sipper counts in early withdrawal (CTRL female = 17.3 ± 3.50, male = 6.5 ± 2.50, OXY female = 62.9 ± 12.33, male = 82.6 ± 20.87, F_drug_ = 24.427, p<0.001). **(f-h)** Both male and female mice that had self-administered OXY exhibited higher sipper counts in early withdrawal (CTRL female = 15.2 ± 5.64, male = 15.9 ± 3.55, OXY female = 59.4 ± 13.58, male = 64.5 ± 8.12, F_drug_ = 29.220, p<0.001).

Our final behavioral test sought to determine whether mice would consume oxycodone despite negative consequences. Negative consequences are frequently modeled by a physical punishment such as foot shock [64–66] or adulteration with aversive tastants, such as quinine [67,68]. Given the analgesic properties of oxycodone that could confound responses to foot shock punishment, we introduced increasing concentrations of quinine to the oxycodone-containing drinking tube, such that the quinine concentration increased by 125 µM every 48 hours (Fig 4A). Both male and female CTRL mice avoided the quinine-containing bottle at the lowest quinine concentration (Fig 4B). By contrast, OXY mice persisted in drinking oxycodone beyond the lowest concentration of quinine, with 2 of 15 mice even drinking oxycodone at the highest concentration tested (Fig 4C, D). There were no sex differences in the AUC of the quinine preference curve (Fig 4E).

**Figure 4.**
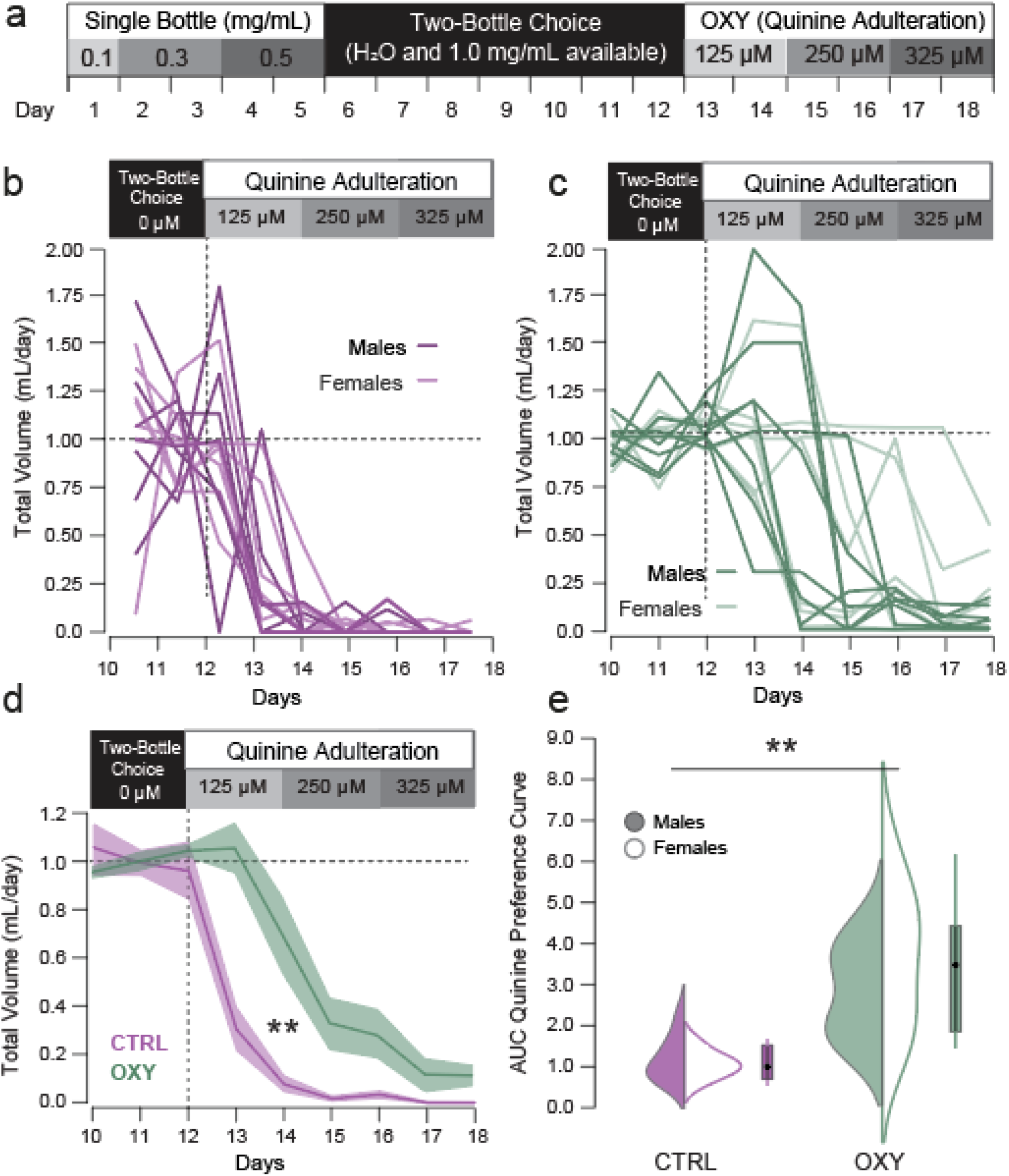
Oral OXY self-administration is resistant to quinine adulteration. **(a)** Experimental schematic. After seven days of two-bottle choice, the OXY-containing bottle (or one H_2_O bottle in CTRL mice) was adulterated with increasing concentrations of quinine. CTRL = 7M/8F, OXY n=8M/7F. **(b-c)** Normalized volume consumed from the quinine-adulterated bottle over the final three days of two bottle choice (before quinine adulteration) and for 6 days of quinine adulteration. **(d-e)** Area under the curve of adulterated quinine drinking volumes. There was a significant effect of Drug (Fdrug = 27.908, p<0.001), of day (F_day_ = 66.767, p<0.001) and interaction (Fdrug●day = 7.485, p<0.001), with both male and female OXY mice exhibiting significantly higher AUC relative to CTRL mice (Fsex = 0.6896, p = 0.414, CTRL female = 0.862 ± 0.130, male = 0.989 ± 0.201, OXY female = 3.495 ± 0.662, male = 2.704 ± 0.415).

### Oxycodone self-administration is characterized by enhanced excitatory transmission in the nucleus accumbens

Addiction is a chronic, relapsing disorder, with behavioral symptoms persisting long into abstinence in humans and rodents [69,70]. The chronic nature of these symptoms is mediated by long-lasting plasticity in the mesolimbic dopamine system, which is induced by exposure to drugs of abuse. One characteristic form of this plasticity that has been causally linked to drug sensitization and persistent relapse is potentiation of excitatory input onto medium sized-spiny neurons in the NAc (MSNs) [71–75]. This potentiation is mediated by the insertion of AMPA receptors into the post-synaptic membrane of accumbal MSNs, is dopamine-dependent, and has been observed after self-administration of opioids [41,76–79] and psychostimulants [72,75,80]. Here, we performed whole cell recordings of accumbal MSNs in the NAc shell after 22 days of withdrawal from oxycodone self-administration (Fig 5A, B).

**Figure 5.**
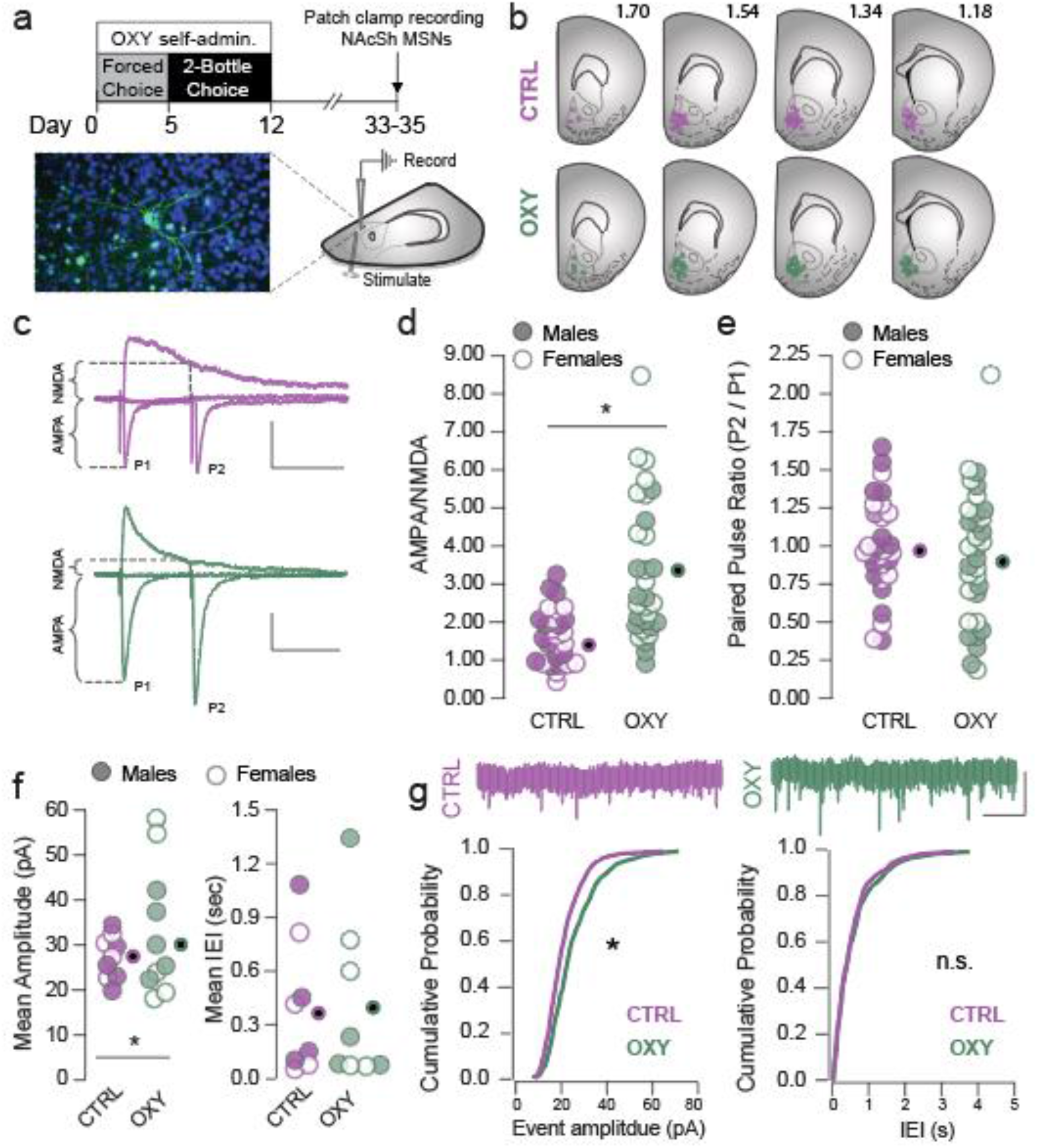
Excitatory transmission onto NAc MSNs is increased after withdrawal from OXY self-administration. **(a)** experimental schematic and representative alexofluor-488 filled MSN (green), DAPI nuclear stain shown in blue. **(b)** Location of recorded cells in control (above) and oxycodone-self-administering groups (bottom); AP coordinates relative to bregma are indicated (mm) **(c)** representative traces showing calculation of AMPA/NMDA and paired pulse ratio in control (top) and OXY withdrawn (bottom) groups; scale bar = 50 pA, 50 ms. **(d)** There was a significant increase in A/N following OXY withdrawal (CTRL = 1.79 ± 0.13, OXY = 3.46 ± 0.33; F_drug_ = 21.07, p<0.0001), and a significant drug by sex interaction (F_drug●sex_ = 4.98, p = 0.030). **(e)** PPR was not different between groups (CTRL = 1.06 ± 0.06, OXY = 0.945 ± 0.07; F_drug_ = 1.37, p=0.36). **(f-g)** There was a significant increase in mEPSC amplitude following OXY withdrawal (Kolmogorov–Smirnov test statistic = 2.89, p < 0.05) but no difference in mEPSC frequency between groups (Kolmogorov–Smirnov test statistic = 1.12, p = 0.16). Scale bar = 20 pA, 1 sec.

The AMPA-to-NMDA ratio (Fig 5C, D) and mEPSC amplitude (Fig 5F) were significantly greater in mice who underwent oxycodone self-administration and withdrawal relative to control mice, suggesting post-synaptic insertion of AMPA receptors. We found no differences in PPR (Fig 5E) nor in frequency of mEPSCs (Fig 5F) between oxycodone self-administering mice and control mice, providing evidence that pre-synaptic excitatory transmission onto accumbal MSNs is not altered by oxycodone self-administration and withdrawal. Together, these results provide evidence that self-administration and withdrawal from oxycodone leads to a post-synaptic potentiation of excitatory synapses onto NAc MSNs, consistent with hallmark plasticity that is induced by self-administration and withdrawal from other opioids and psychostimulants.

## Discussion

Much of our current understanding of addiction has relied on animal models to understand the relationship between behavioral phenomena and their neurobiological correlates. Environmental factors, route of administration, and pattern of intake strongly influence nearly every aspect of drug misuse, yet previous models of addiction have historically been conducted under experimental parameters which differ significantly from typical conditions associated with prescription opioid misuse. For example, intermittent access to i.v. drug infusions with discrete cues in a novel environment are ideal for modeling addiction to psychostimulants. However, prescription opioids, including oxycodone, are typically administered via the oral route, with near continuous access (especially with extended-release formulations) in familiar environments. Here, we characterize a novel paradigm that more closely recapitulates factors relevant to prescription opioid abuse, which may prove valuable for studying the neurobiology of opioid use disorder in mice.

In this oral oxycodone self-administration paradigm, female mice consumed more oxycodone (mg/day) during the single bottle phase and higher total dose of oxycodone (mg/kg) compared to males (Fig 1), consistent with previous observations that female rodents self-administer higher doses of opioids than males [3,5,81–84]. The effects of sex on opioid intake, dependence, and abuse liability have been well documented [61,81,85–88], and interact with feeding conditions [89,90] and route of administration [3,38] to influence the bioavailability of oxycodone. While pharmacokinetic influences on sex differences were beyond the scope of our study, others have explored sex-dependent factors that may contribute to different oral oxycodone consumption between females and males. Phillips et al. (2018) [3] demonstrate that while oral oxycodone ubiquitously resulted in elevated plasma concentrations of oxycodone and its active metabolite oxymorphone, the extent of this effect was influenced by sex, presence of sex organs, and feeding conditions, and was exacerbated following repeated administrations. Such observations are congruent with pharmacokinetic studies showing that plasma concentrations of oxycodone and its active metabolites are mediated, in part, by CYP3A4 and CYP2D6 [91,92], whose activity can be modulated by sex hormones and food intake [93,94]. Importantly, however, sex effects on oxycodone pharmacokinetics were either diminished or not observed with i.v. administration [38,95], highlighting the importance of route of administration in addiction models.

Moreover, mice voluntarily escalate their consumption of oxycodone despite ad libitum access to drinking water (Fig 1, S1), and alter their pattern of intake (Fig S2, S3). Escalation of intake is a key feature which is relevant to modeling DSM criteria for addiction [11,96,97]. Additionally, we found that voluntary oxycodone consumption induced robust physical withdrawal signs, including hyperactivity, jumps and paw tremors which persisted into protracted withdrawal, albeit with reduced severity than in acute withdrawal (Fig 2). While there was large variability in the severity of observable withdrawal signs, it is critical that this paradigm fomented these signs; withdrawal symptoms often contribute to drug relapse through negative reinforcement, where the drug is taken to relieve the aversive state of withdrawal [98,99].

Another cardinal DSM criterion for OUD is drug craving [40], which is frequently modeled with reinstatement paradigms. In these classical reinstatement paradigms, an instrumental response is first paired with drug infusion in a self-administration paradigm. Then, following a period of extinction training, stress, a priming dose of drug, or discriminative or contextual cues are introduced to reinstate the instrumental response in the absence of further drug availability [10,20,101]. However, this extinction training is not typically a feature of prescription drug use. Rather, abstinence is typically imposed by the cessation of opioid availability, either because of abrupt changes in prescribing practices, admission to in-patient facilities, or loss of access to recreational sources [102]. For this reason, we opted to use a ‘forced abstinence’ model, where no extinction training occurs after drug self-administration, and the response that previously lead to drug consumption is measured after variable periods of withdrawal [62,103]. We found that lickometer interactions, which resulted in the delivery of oxycodone solution during the self-administration phase, were significantly greater in oxycodone-self-administering mice relative to controls at both acute (24h) and late (3 week) withdrawal phases (Fig 3). Interestingly, we did not find evidence for the incubation of craving in this paradigm, which refers to the tendency for craving-related behavior to increase with the length of the forced abstinence period [62,122]. We propose three possible explanations for this: incubation of craving is most robustly observed under long access, high concentrations of IV administration, where initial seeking responses on the first probe session are typically low [72]. This is not the case with our oral administration, where seeking behavior is quite high, even during a short assay in the acute withdrawal. Additionally, incubation of craving is frequently measured when mice are tested either in acute or late withdrawal, and behavior is compared between different subjects, with late withdrawal subjects showing increased seeking [75,104]. Here, we are testing the same subjects at two time points, and while we don’t see differences between these time points, it is possible that the first session served as an extinction trial, which may mask any potential effects on incubation [105]. Finally, the time course of incubation of craving is highly dependent on the addictive substance [62], and the single three-week time point may not be adequate to capture the temporal dynamics of oxycodone seeking after withdrawal.

We modeled drug consumption despite negative consequences by adulterating oxycodone solution with increasing doses of quinine, which is commonly used to assess compulsive alcohol drinking [106,107]. While the quinine concentrations were quite low relative to ethanol studies, oxycodone is an alkaloid with a bitter taste prior to the introduction of quinine and already aversive, which we confirmed by establishing that naïve mice avoid 1.0 mg/mL oxycodone before the introduction of quinine (Fig S1A). Critically, we also observed individual differences in quinine-resistant drug consumption, with only a subset of mice persisting at the highest quinine doses (Fig 4). This is reminiscent of i.v. opioid or psychostimulant self-administration, where typically only a subpopulation (15-30%) of animals exhibit punishment-resistant drug self-administration [65,108–110]. This is critical as it also reflects human patients who are often able to use drugs recreationally, with only a subpopulation transitioning to compulsive drug use [109,111] and underscores the utility of this model for studying individual differences in aversion-resistant oxycodone consumption.

Finally, we demonstrate that oral oxycodone self-administration induces potentiation of excitatory transmission into NAc MSNs (Fig 5), which has been linked to persistence of reinstatement and physical symptoms following withdrawal from self-administration of cocaine [42,75,112] and opioids [113–117]. This plasticity is induced by drug-induced dopamine release from the ventral tegmental area (VTA) into the nucleus accumbens [42,43,118], and can be mimicked by strong optogenetic stimulation of VTA DA neurons alone [65]. Relative to morphine, oxycodone causes even more robust release of dopamine into the NAc, presumably through mu-receptor activation of inhibitory inputs to VTA dopamine neurons [119]. Consistent with this mechanism, we also found that oxycodone self-administering mice exhibited an increase in AMPA/NMDA ratio in accumbal MSNs, which was accompanied by an increased amplitude of mEPSCs. Conversely, we observed no effect of oxycodone self-administration on PPR or frequency of mEPSCs. Together, these data suggest a post-synaptic strengthening of excitatory transmission onto accumbal MSNs. Cocaine exposure potentiates excitatory transmission onto NAcSh MSNs, although this plasticity is preferentially expressed in D1-MSNs [65,121,123] (but see [72]), while morphine self-administration potentiates excitatory drive onto both D1- and D2-MNSs [41,78]. Here, we used electrical stimulation in wild-type mice, and therefore do not know if adaptations we observe are specific to a particular excitatory accumbal input, or whether they occur in both D1- and D2-MSNs within the accumbens. However, our results suggest a bimodal distribution of AMPA/NMDA ratio, raising the possibility that adaptations may be cell-type specific. Moreover, the paradigm of drug access differentially affects plasticity at discrete excitatory inputs to the NAc, with inputs from the paraventricular thalamus implicated in physical signs of withdrawal [77,78], medial prefrontal cortical inputs implicated in drug seeking, sensitization and reinstatement [41,72,79,120,121], and hippocampal inputs implicated in context recognition necessary for behavioral expression of drug seeking [73]. While these results demonstrate the utility of this oral paradigm for studying synaptic mechanisms underlying persistent drug-adaptive behavior into withdrawal, future work will determine how pathway- and cell type-specific plasticity contribute to discrete elements of addiction-relevant behavior in this oral opioid self-administration paradigm.

Here we characterize a novel paradigm for study prescription opioid misuse. This oral, home cage two-bottle choice procedure can be used to model several features of DSM criteria that are specifically relevant to OUD, while conserving the route of administration, pattern of access, and contextual factors relevant to prescription opioid intake. This model has potential utility for studying the neurobiological substrates driving drug relapse and quinine-resistant drug intake. Moreover, the high-throughput nature of this paradigm permits high-powered studies necessary for understanding sex differences, circadian biology, and individual variability underlying vulnerability to OUD.

## Supporting information

Combined Supplemental Figures

## Acknowledgements

We thank M.C. Stander and E. Godynyuk for excellent technical help. We also J.J. Choong, S. Golden and M. Mathis for their commitment to open-source tools and for support with supervised behavioral classification (SimBA) and markerless pose estimation (DeepLabCut^TM^), respectively. We thank MCCI Corporation (Ithaca NY) for development of wireless in-cage activity and environmental sensors and setting up the IoT cloud solution.

## Author Contributions

CM, YHC, RP and YMV performed behavioral experiments. CM, JL, JRT, BC and MCC performed patch-clamp experiments. AVK and MCC developed home cage-based lickometer devices for self-administration, TE developed custom software for analysis. CM, BC, and MCC analyzed data. RG, BC, AVK, and MCC supervised work. CM, YHC, and MCC wrote the manuscript with input from all authors.

---

**Supplemental Figure 1.**
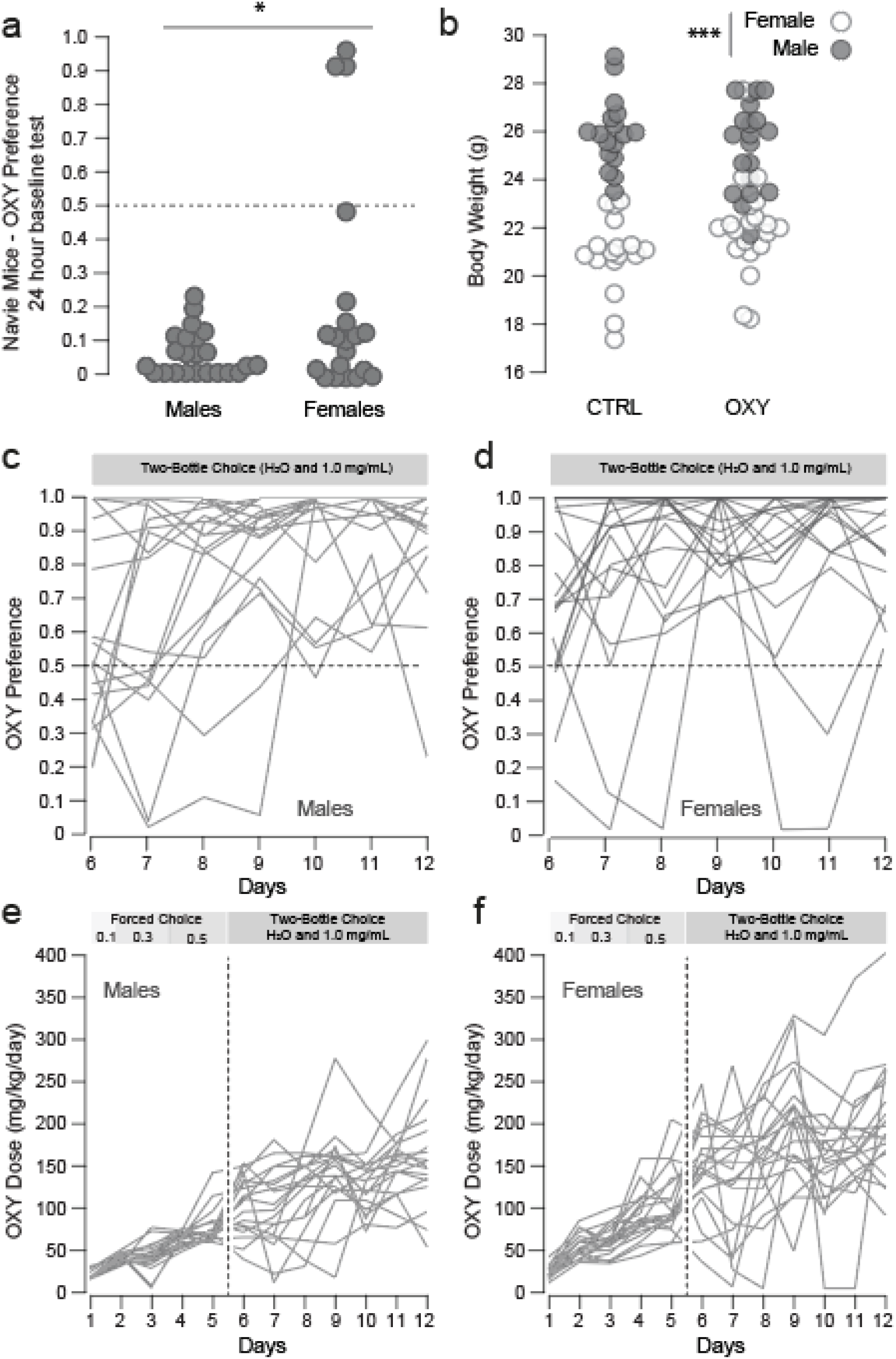
Individual data for oxycodone dose and preference. **a)** Baseline preference for 1.0 mg/ml oxycodone in naïve mice; while both males and females showed a preference for water over oxycodone, the initial preference was higher for females, which was driven by a subset of oxy-preferring mice (males: 5.21 ± 1.39, females: 24.03 ± 7.90, F_sex_ = 6.59, p = 0.014). **b)** Body weight for individual subjects; while the average weight of males was significantly higher than that of females (CTRL male = 26.09 ± 37, female = 20.96 ±0.39, OXY male = 25.57 ± 0.41, female = 22.07 ± 0.46), drug treatment did not have a significant effect on body weight (F_sex_ = 107.174, p< 0.0001, F_drug_ = 0.474, p = 0.493, F_sex*drug_ = 3.837, p = 0.054). (c-d) Daily dose escalation curves for individual subjects (n = 20 M, 20 F). **(c-d)** Preference for oxycodone for individual mice; preference is defined as [*consumed volume of oxycodone / total volume of fluid consumption*]. **(e-f)** Escalation of oxycodone dose for individual mice; dose is defined as [*mg oxycodone consumed / body weight*].

**Supplemental Figure 2.**
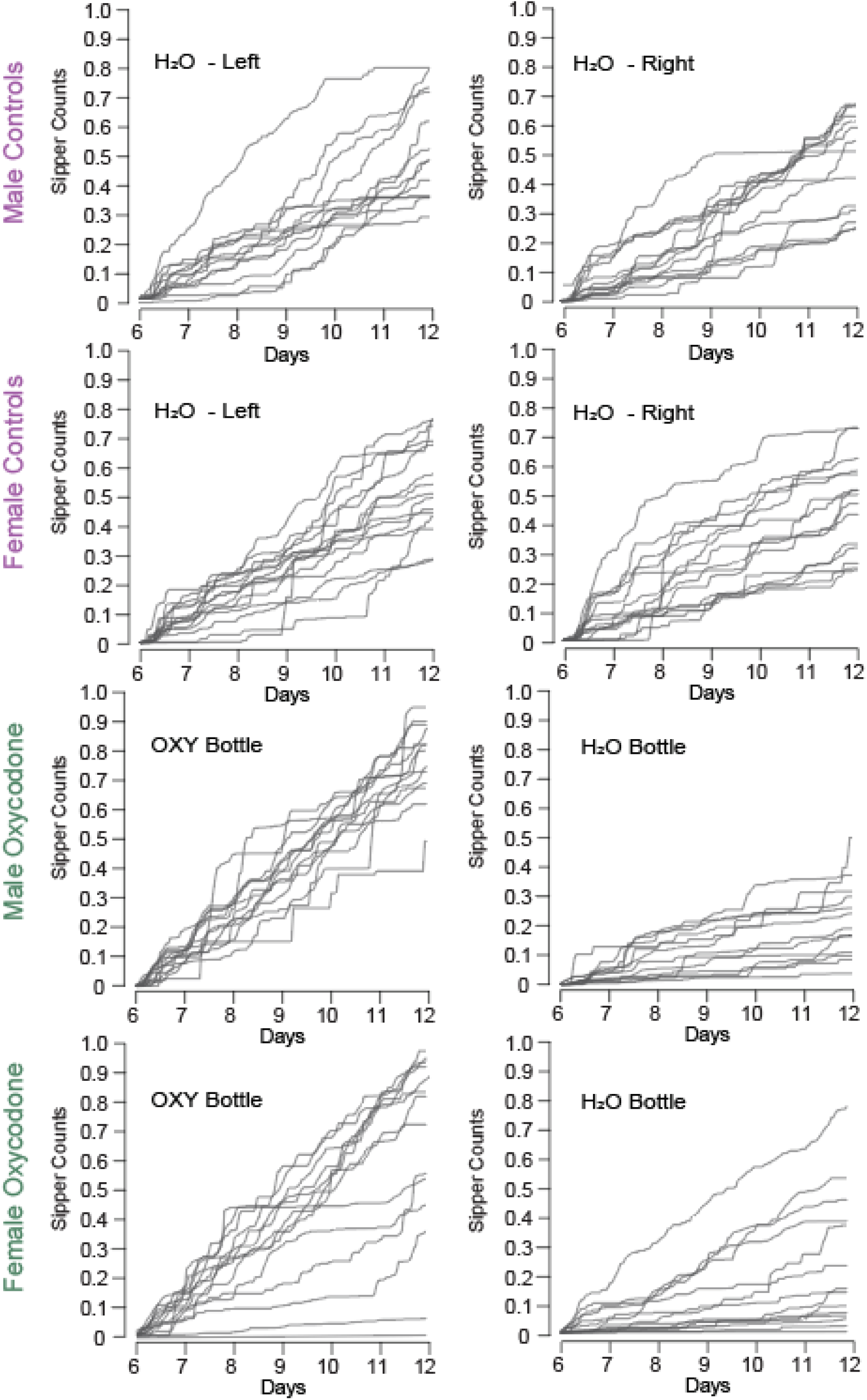
Individual data for sipper counts over two-bottle choice phase in control and oxycodone self-administering mice. Individual cumulative lickometer photo-interrupter counts for each subject are shown. Counts made on the left and right bottles are shown for controls are shown separately (left, right, respectively). Counts made on the oxycodone-filled and water-filled bottles are shown separately (oxycodone, water, respectively).

**Supplemental Figure 3.**
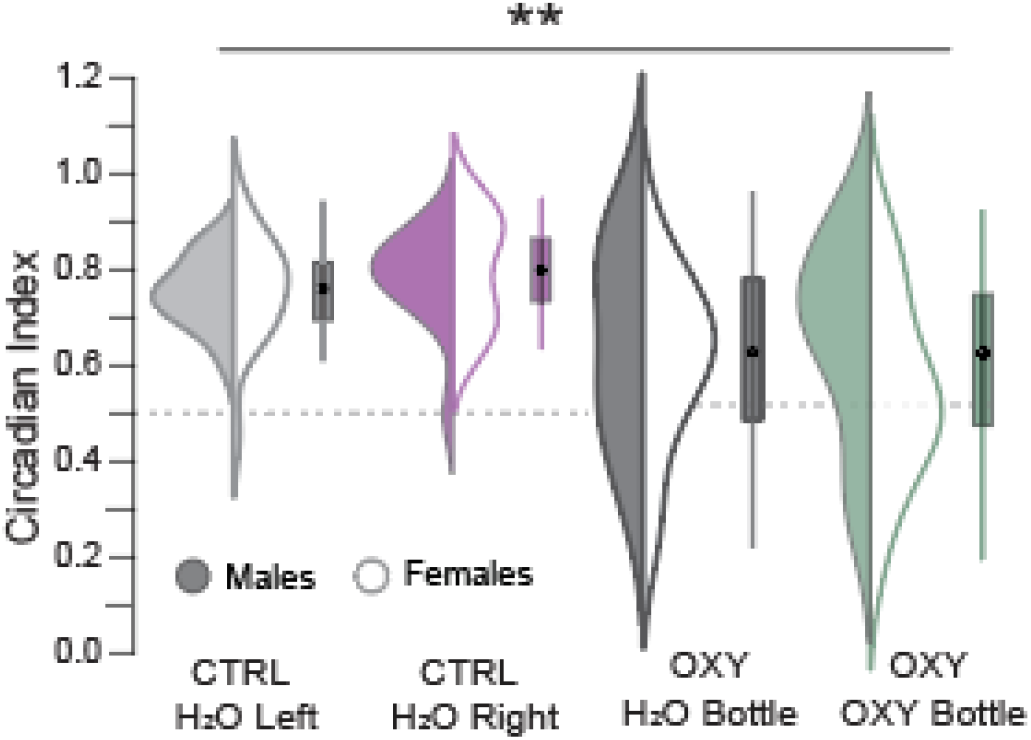
Homecage drinking monitoring confirms OXY preference and reveals altered patterns of OXY and water intake. Circadian index for Sipper Counts on each bottle (Dark Cycle Counts/Light Cycle Counts). Circadian index for both water and oxycodone drinking was significantly lower in OXY-mice (CTRL female Left/Right = 0.740 ± 0.030/0.783 ± 0.029, male Left/Right = 0.737 ± 0.021/0.768 ± 0.029, F_Drug_ = 28.617 p < 0.001, F_Bottle_ = 0.899, p = 0.345).

**Supplemental Figure 4.**
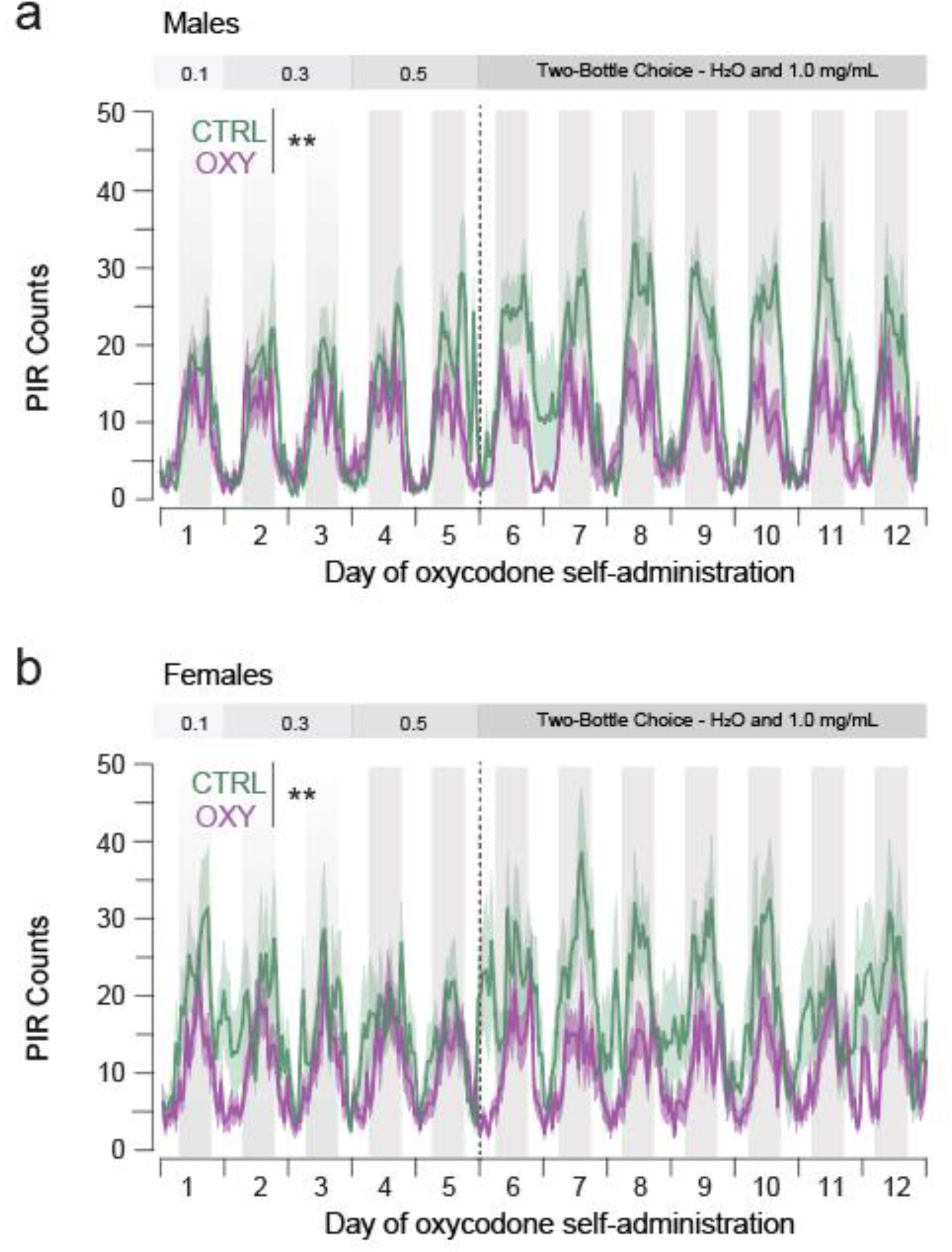
Oxycodone-induced activity patterns did not differ between male and female mice. PIR counts during the single bottle and two-bottle choice phases of the self-administration protocol are shown for male mice **(a)** and female mice **(b)**. **P < 0.01

**Supplemental Figure 5.**
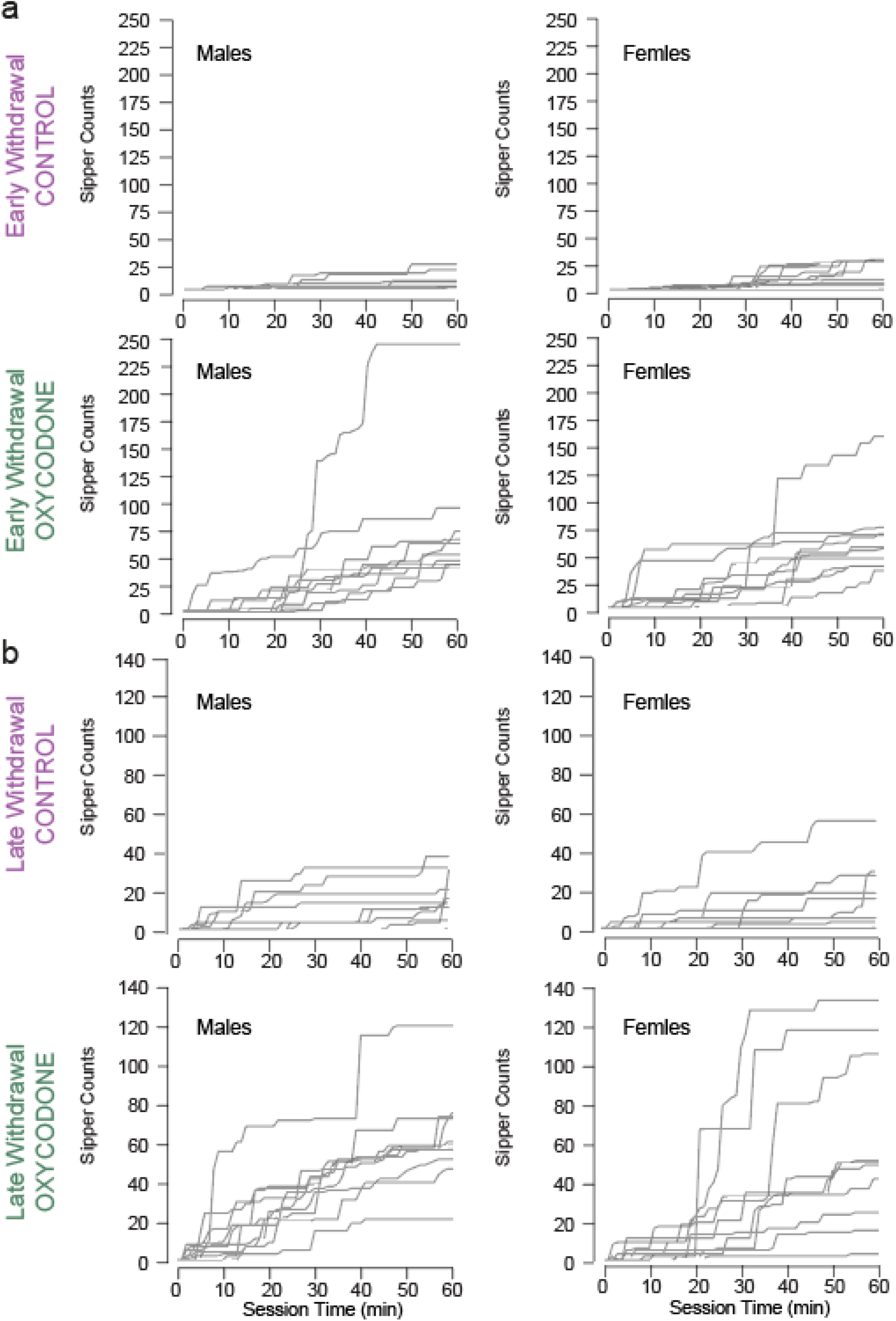
Individual data for sipper counts during early and late seeking tasks. Individual photobeam breaks made during the 60-minute probe test made under extinction conditions in both the early **(a)** and late **(b)** withdrawal time points. Data from males (left) and females (right) is shown.

