## Supplementary material for "Modeling features of addiction with an oral oxycodone self-administration paradigm": Combined Supplemental Figures

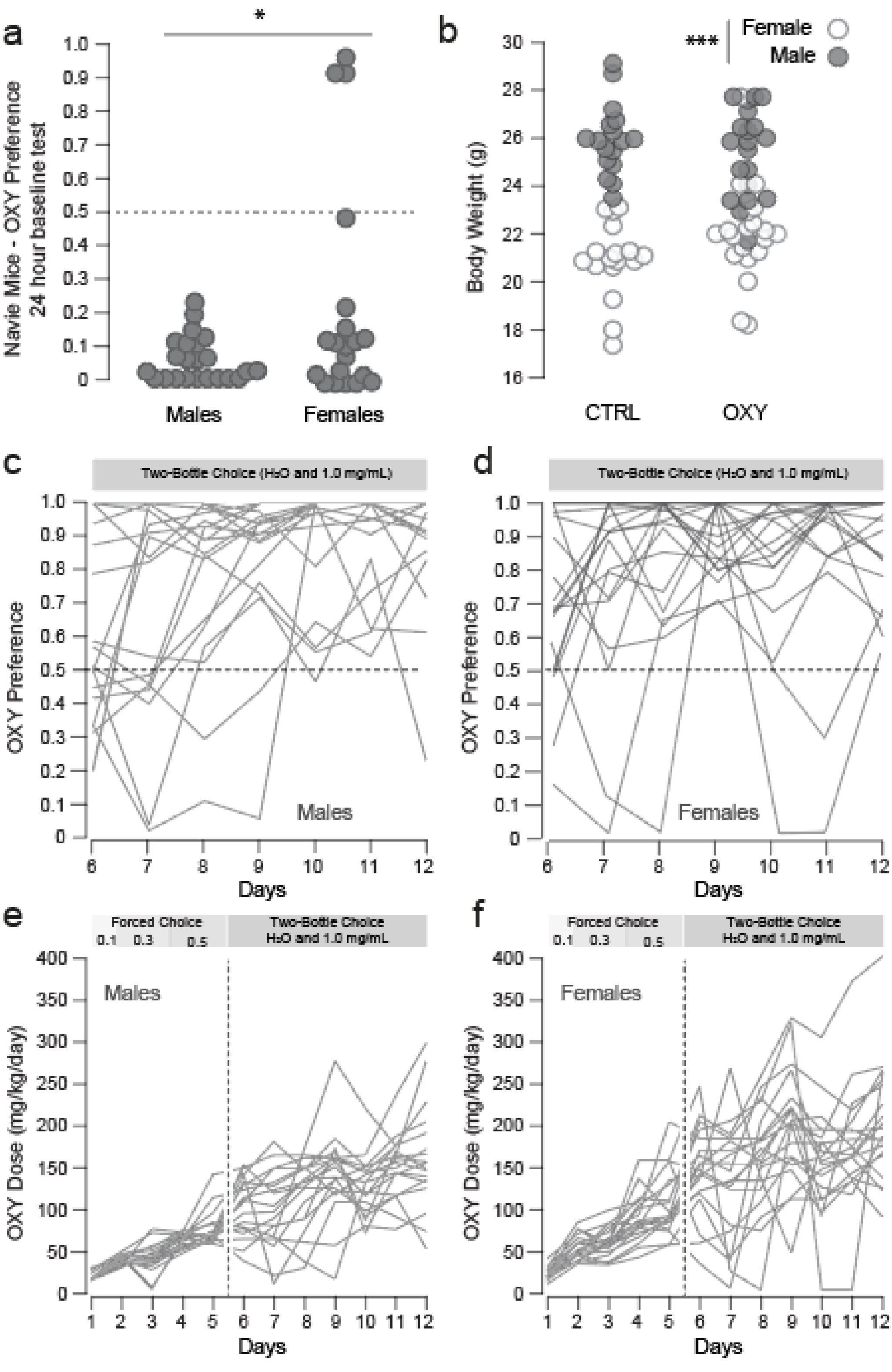


**Supplemental Figure 1. Individual data for oxycodone dose and preference.**

**Supplemental Figure 1. Individual data for oxycodone dose and preference. a)** Baseline preference for 1.0 mg/ml oxycodone in naïve mice; while both males and females showed a preference for water over oxycodone, the initial preference was higher for females, which was driven by a subset of oxy-preferring mice (males: 5.21 ± 1.39, females: 24.03 ± 7.90, $F_{sex}$ = 6.59, p = 0.014). **b)** Body weight for individual subjects; while the average weight of males was significantly higher than that of females (CTRL male = 26.09 ± 37, female = 20.96 ±0.39, OXY male = 25.57 ± 0.41, female = 22.07 ± 0.46), drug treatment did not have a significant effect on body weight ($F_{sex}$ = 107.174, p< 0.0001, $F_{Drug}$ = 0.474, p = 0.493, $F_{sex*drug}$ = 3.837, p = 0.054). (c-d) Daily dose escalation curves for individual subjects (n = 20 M, 20 F). **(c-d)** Preference for oxycodone for individual mice; preference is defined as [*consumed volume of oxycodone / total volume of fluid consumption*]. **(e-f)** Escalation of oxycodone dose for individual mice; dose is defined as [*mg oxycodone consumed / body weight*].

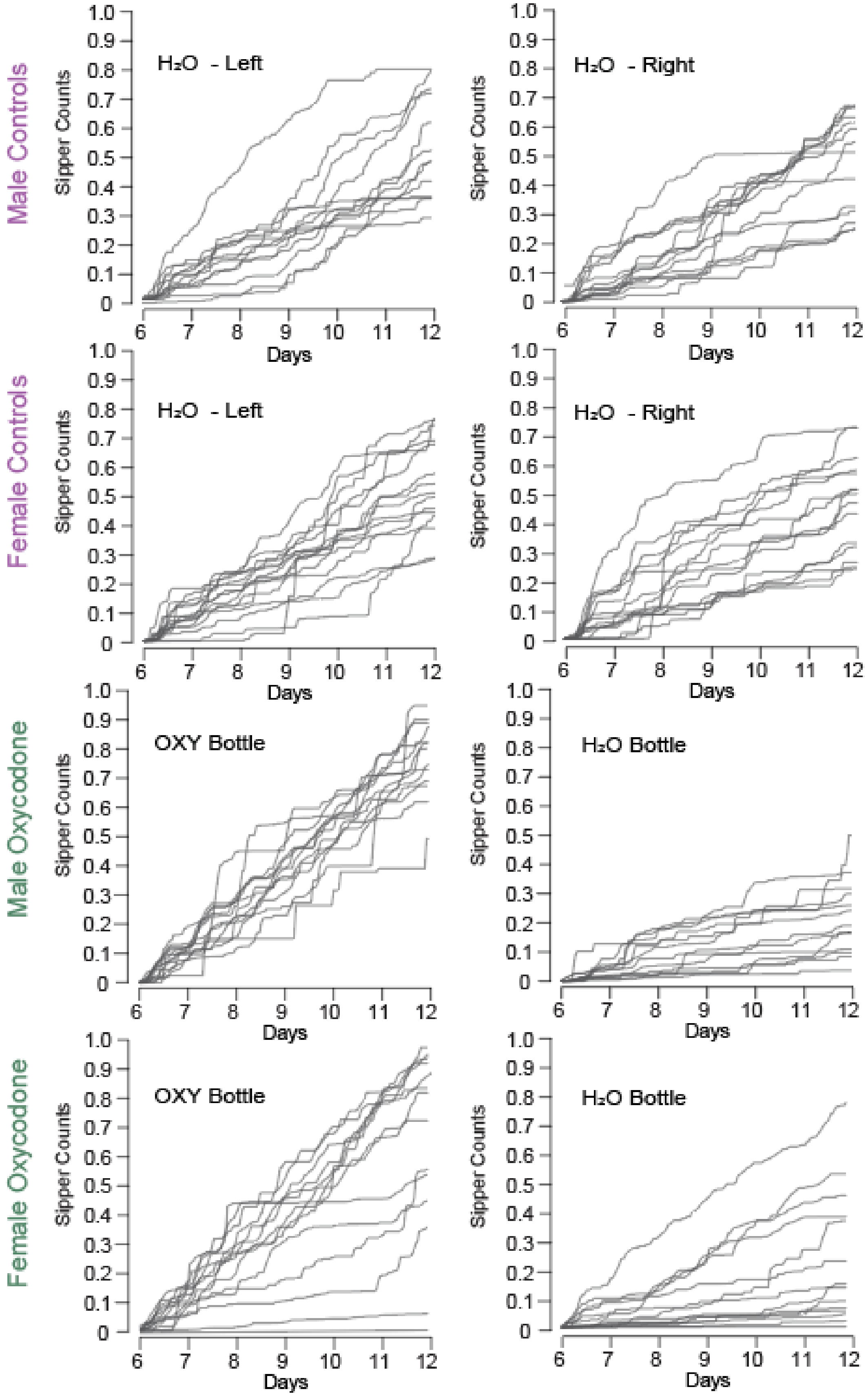


**Supplemental Figure 2. Individual data for lickometer counts during two-bottle choice phase in control and oxycodone self-administering mice.**

**Supplemental Figure 2. Individual data for sipper counts over two-bottle choice phase in control and oxycodone self-administering mice.** Individual cumulative lickometer photo-interrupter counts for each subject are shown. Counts made on the left and right bottles are shown for controls are shown separately (left, right, respectively). Counts made on the oxycodone-filled and water-filled bottles are shown separately (oxycodone, water, respectively).

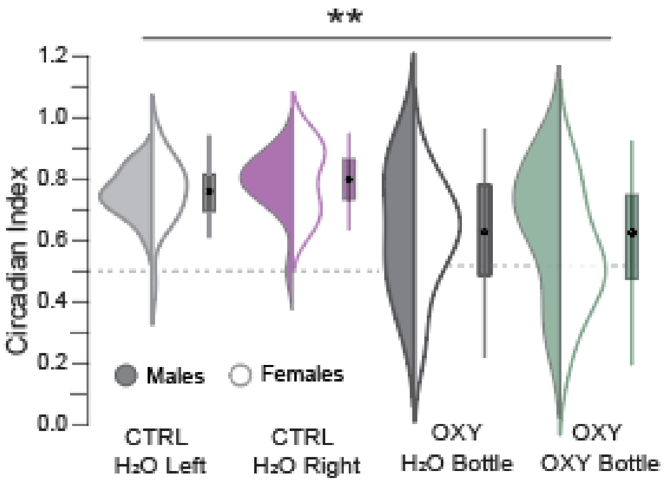


**Supplemental Figure 3. Home cage drinking monitoring confirms altered circadian rhythm of oxycodone intake.**

**Supplemental Figure 3. Home cage drinking monitoring confirms altered circadian rhythm of oxycodone intake.** Circadian index for Sipper Counts on each bottle (Dark Cycle Counts/Light Cycle Counts). Circadian index for both water and oxycodone drinking was significantly lower in OXY-self administering mice (CTRL female Left/Right = 0.740 ± 0.030/0.783 ± 0.029, male Left/Right = 0.737 ± 0.021/0.768 ± 0.029, $F_{Drug} = 28.617$ $p < 0.001$, $F_{Bottle} = 0.899$, $p = 0.345$).

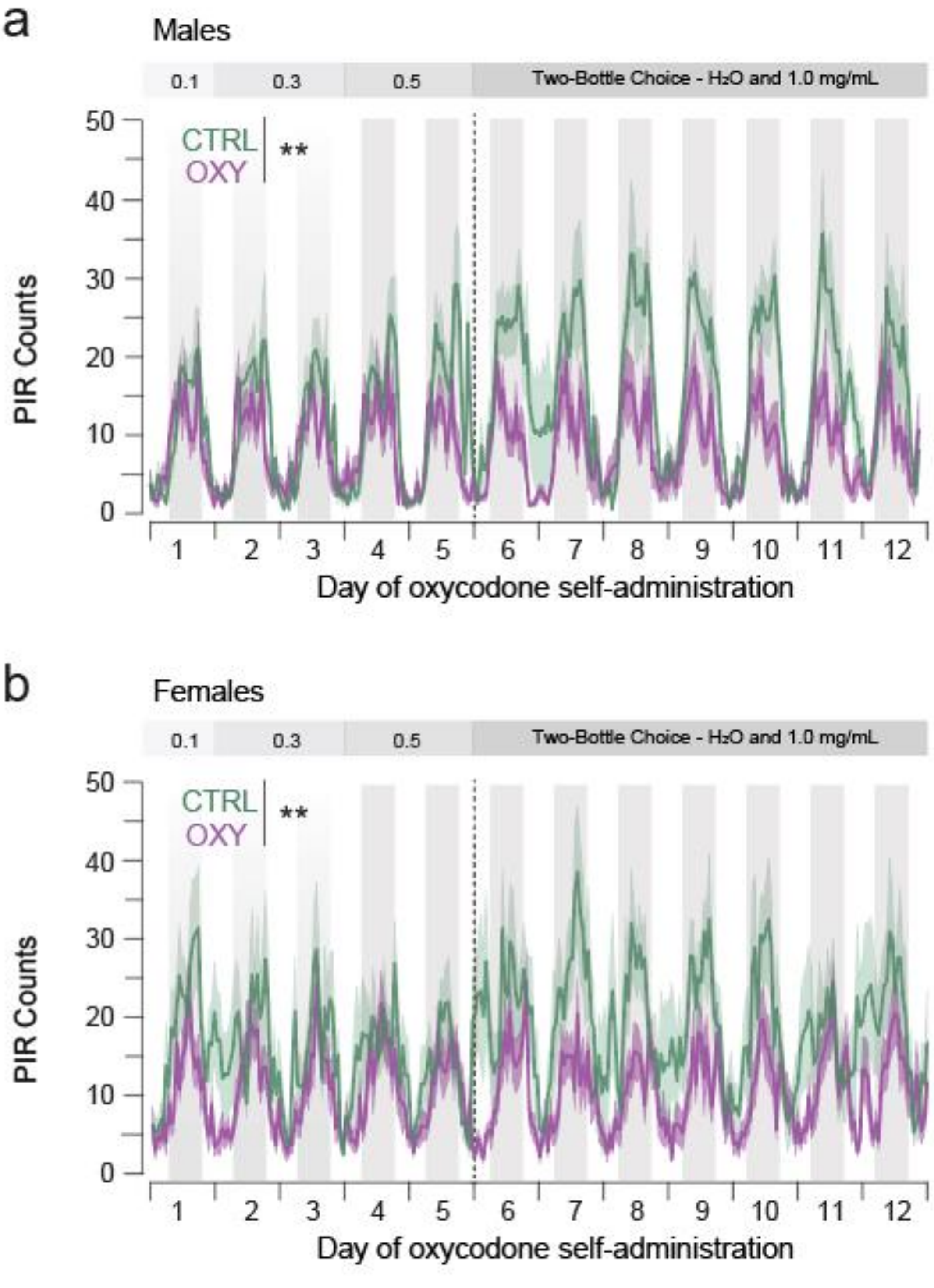


**Supplemental Figure 4. Oxycodone induced activity patterns did not differ between male and female mice.**

**Supplemental Figure 4. Oxycodone-induced activity patterns did not differ between male and female mice.** PIR counts during the single bottle and two-bottle choice phases of the self-administration protocol are shown for male mice **(a)** and female mice **(b)**. $^{**}P < 0.01$

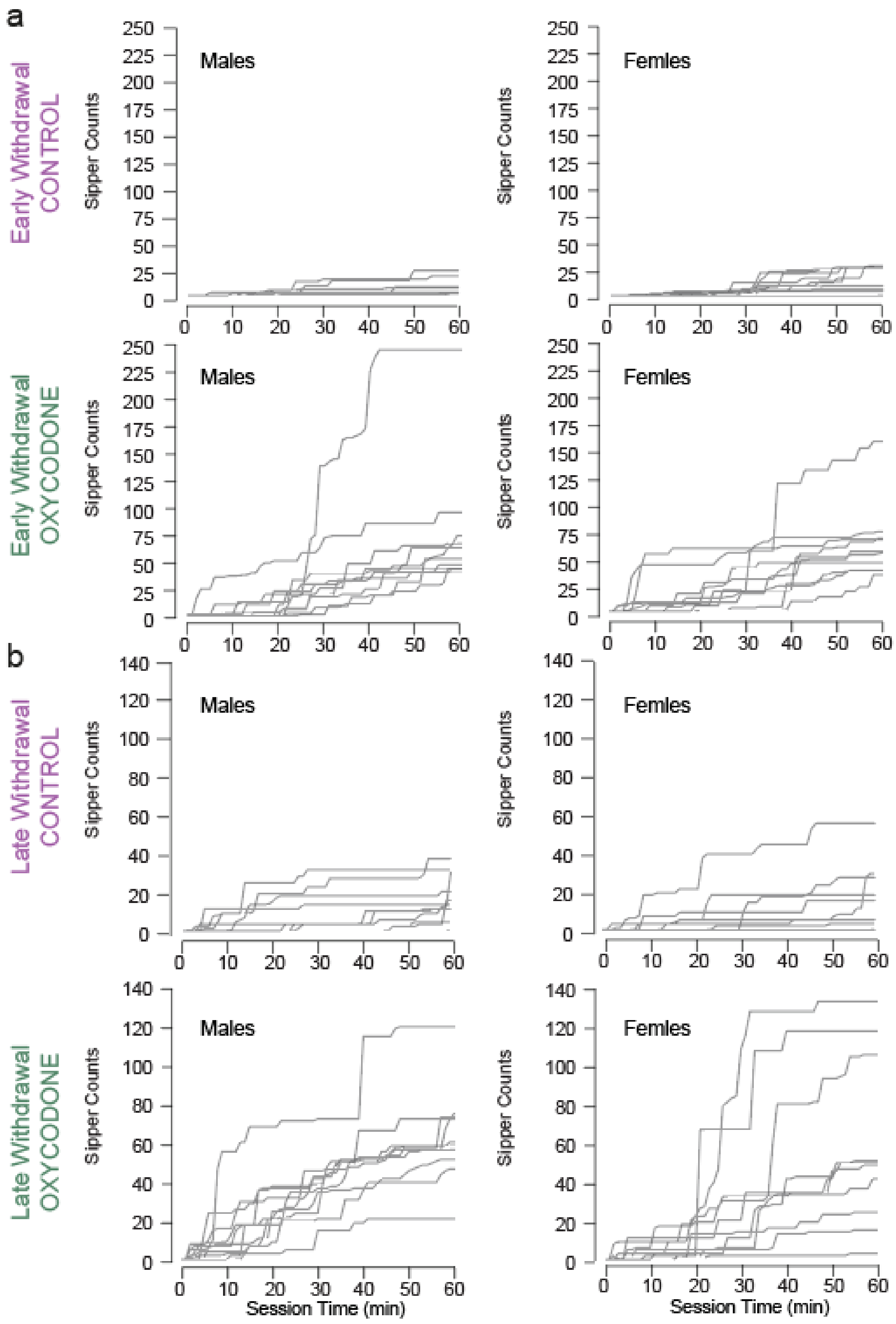


**Supplemental Figure 5. Individual data for oxycodone seeking in extinction after acute and protracted withdrawal.**

**Supplemental Figure 5. Individual data for sipper counts during early and late seeking tasks.** Individual photobeam breaks made during the 60 minute probe test made under extinction conditions in both the early **(a)** and late **(b)** withdrawal time points. Data from males (left) and females (right) is shown.
